## Supplementary figures and images for "AI-directed voxel extraction and volume EM identify intrusions as sites of mitochondrial contact"

### Supplemental Figures

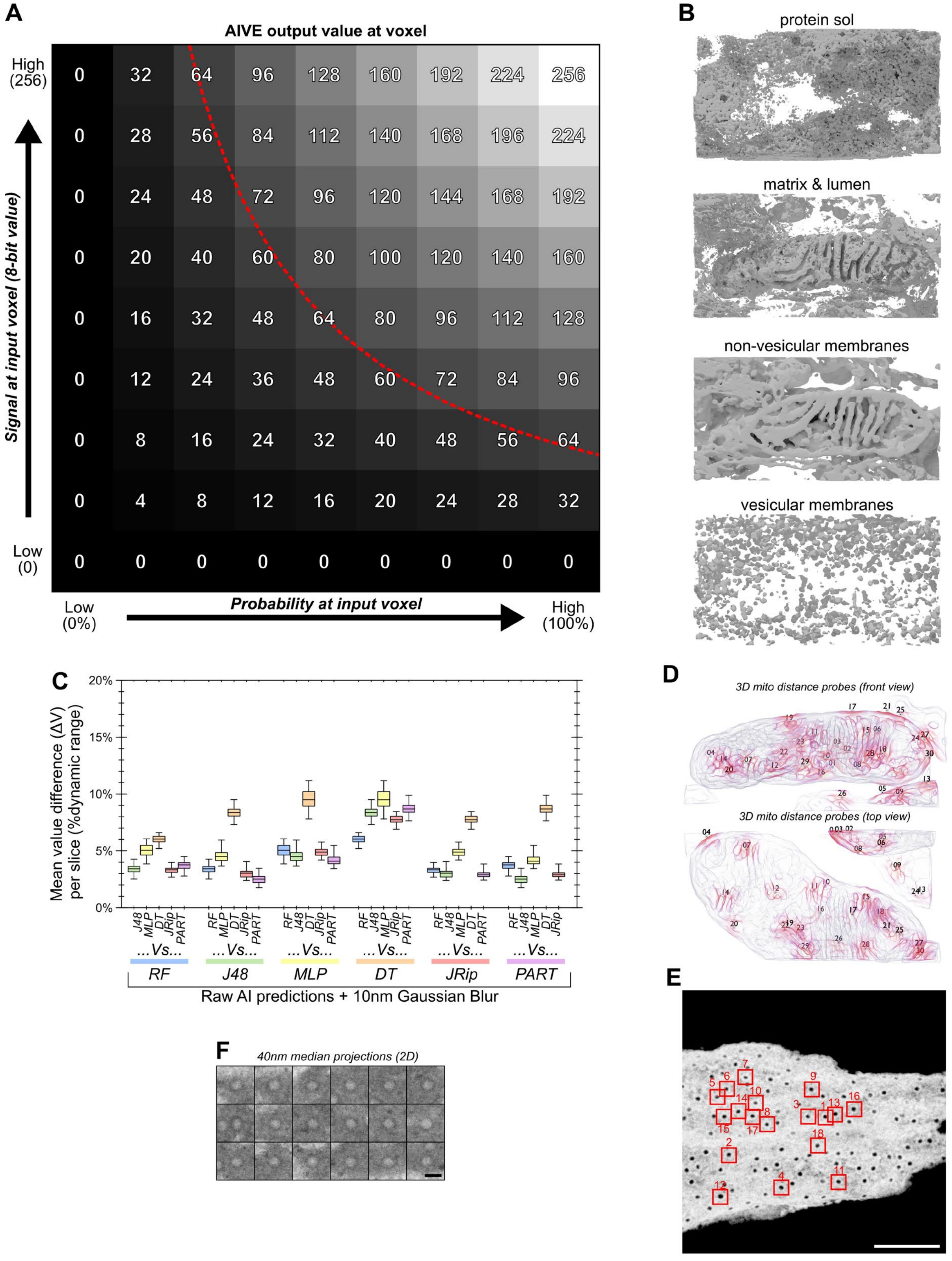

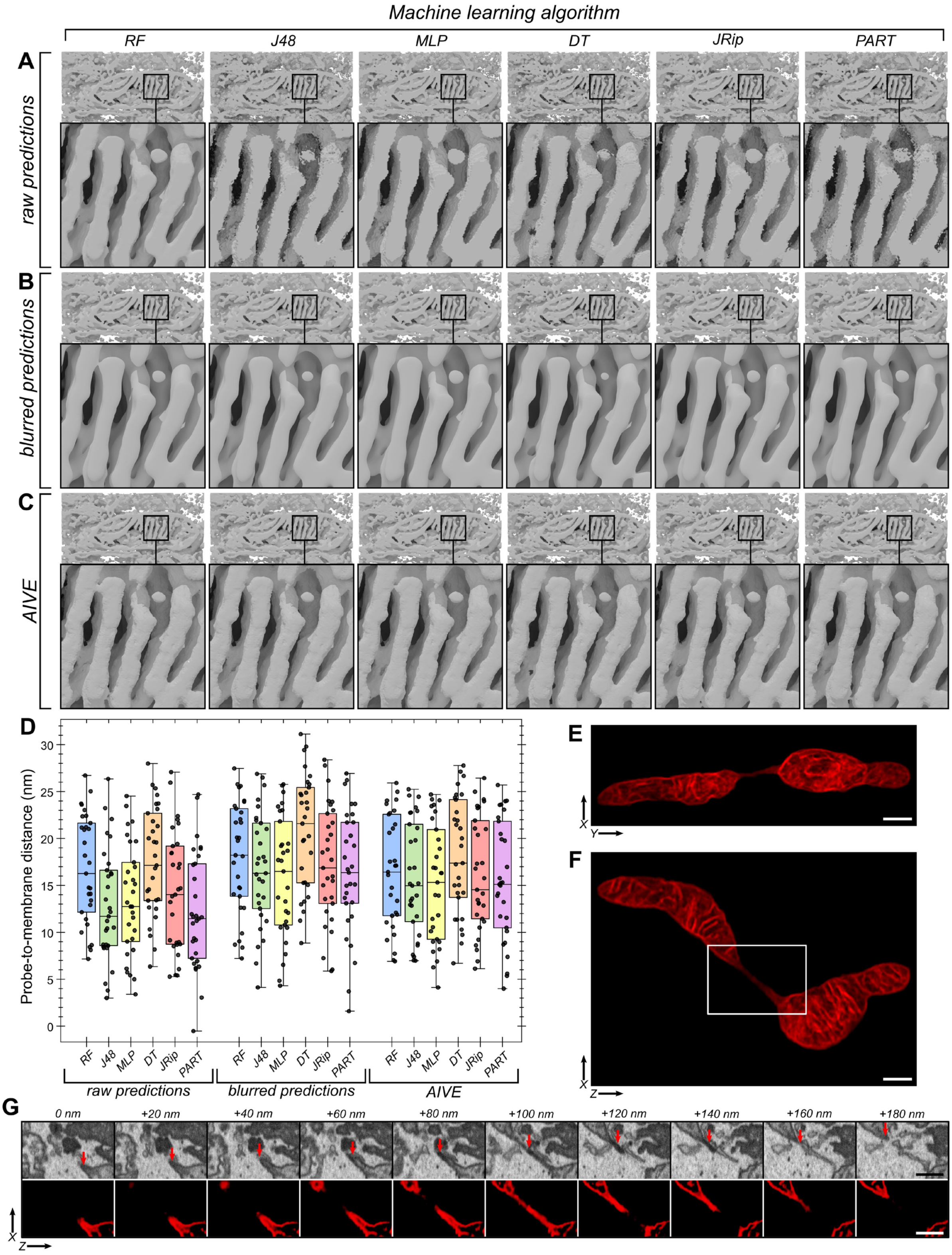

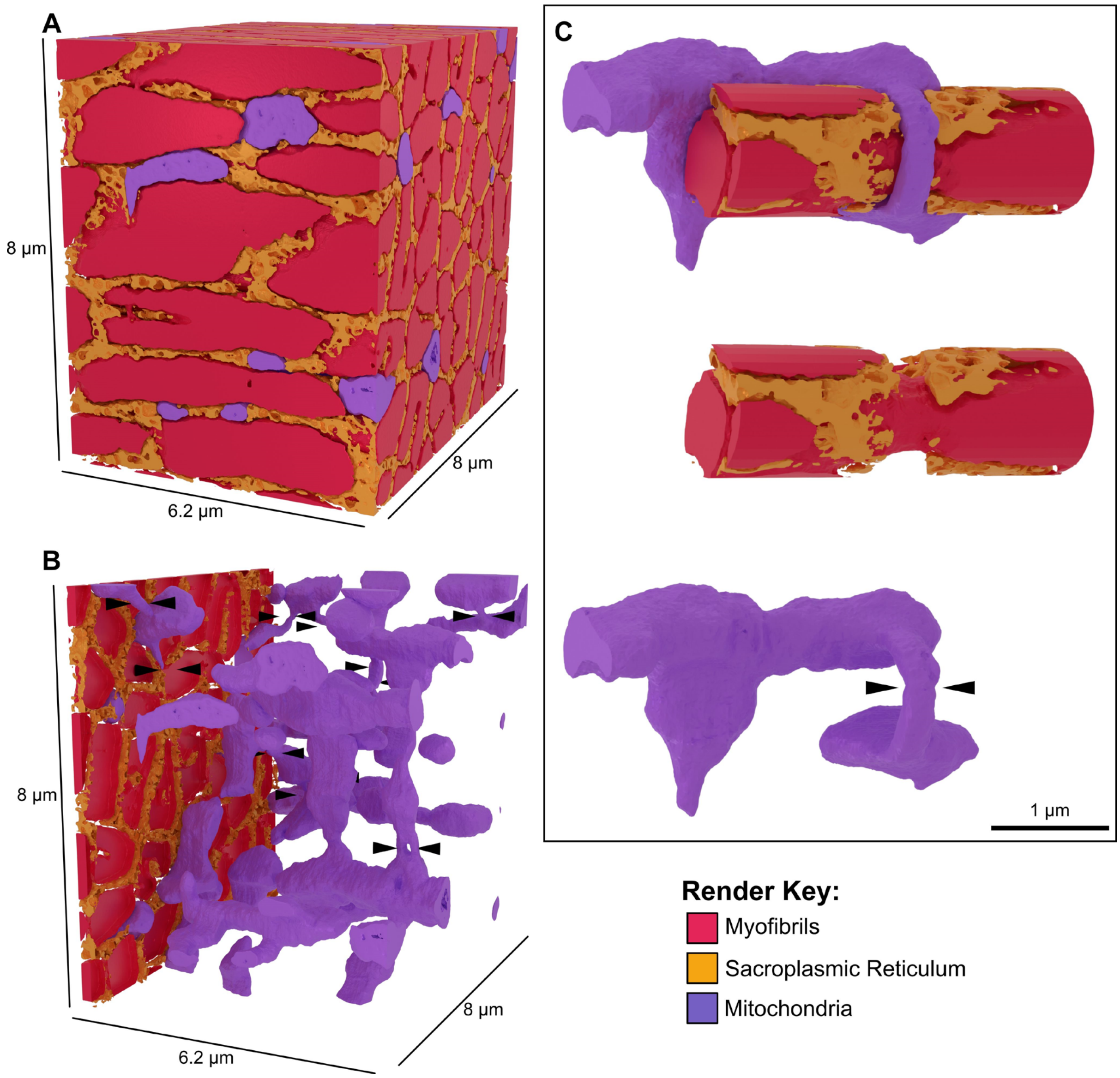

#### Figure S4

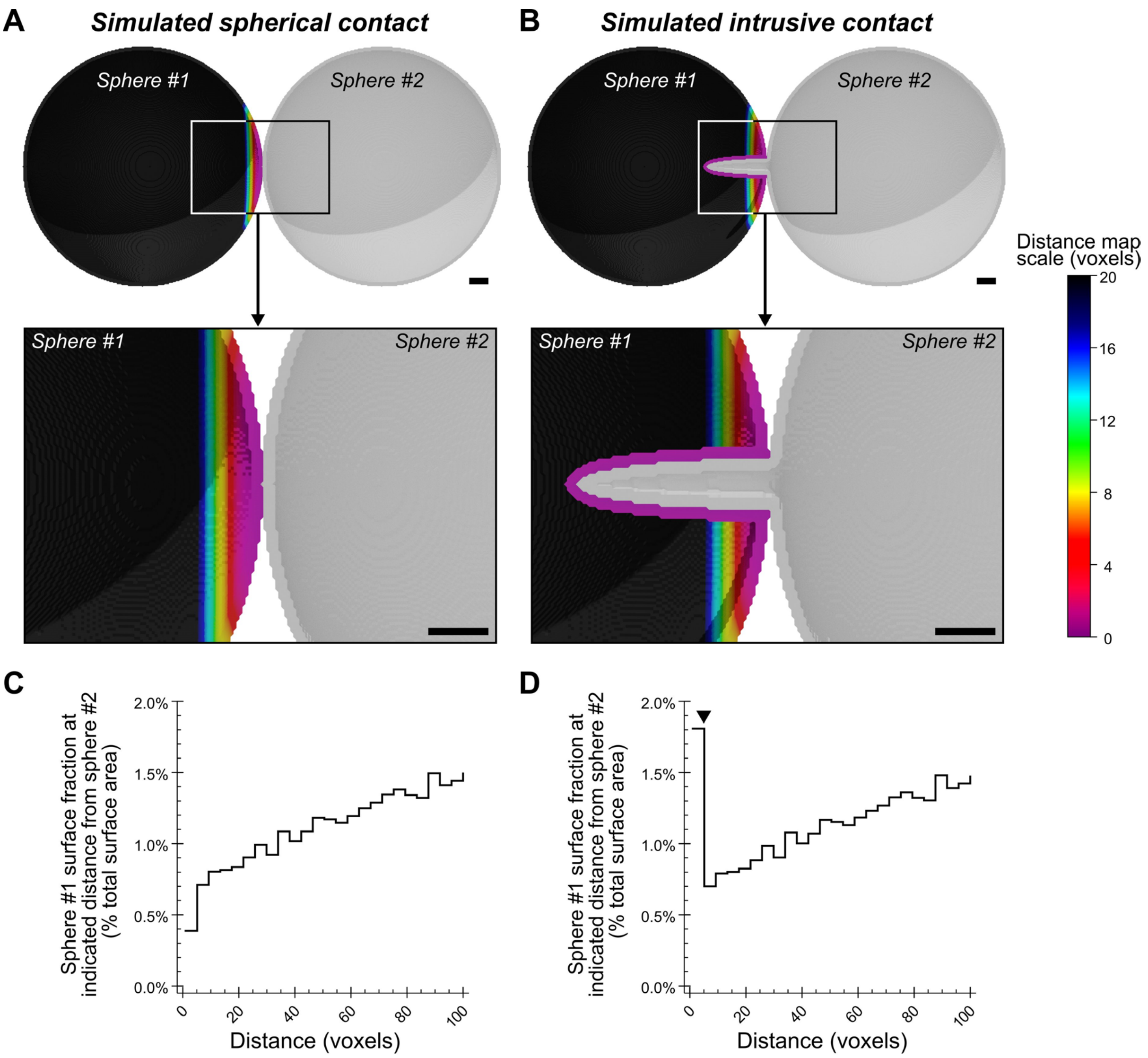


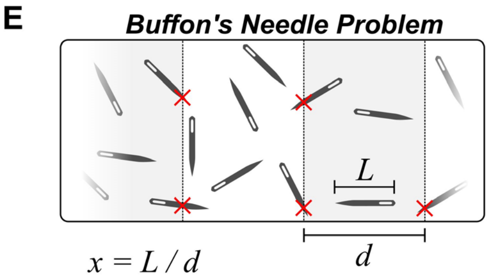


$$P(x) = \begin{cases} \frac{2x}{\pi} & \text{if } x \leq 1 \\ \frac{2}{\pi}\left(x - \sqrt{x^2-1} + \sec^{-1}(x)\right) & \text{if } x > 1 \end{cases}$$

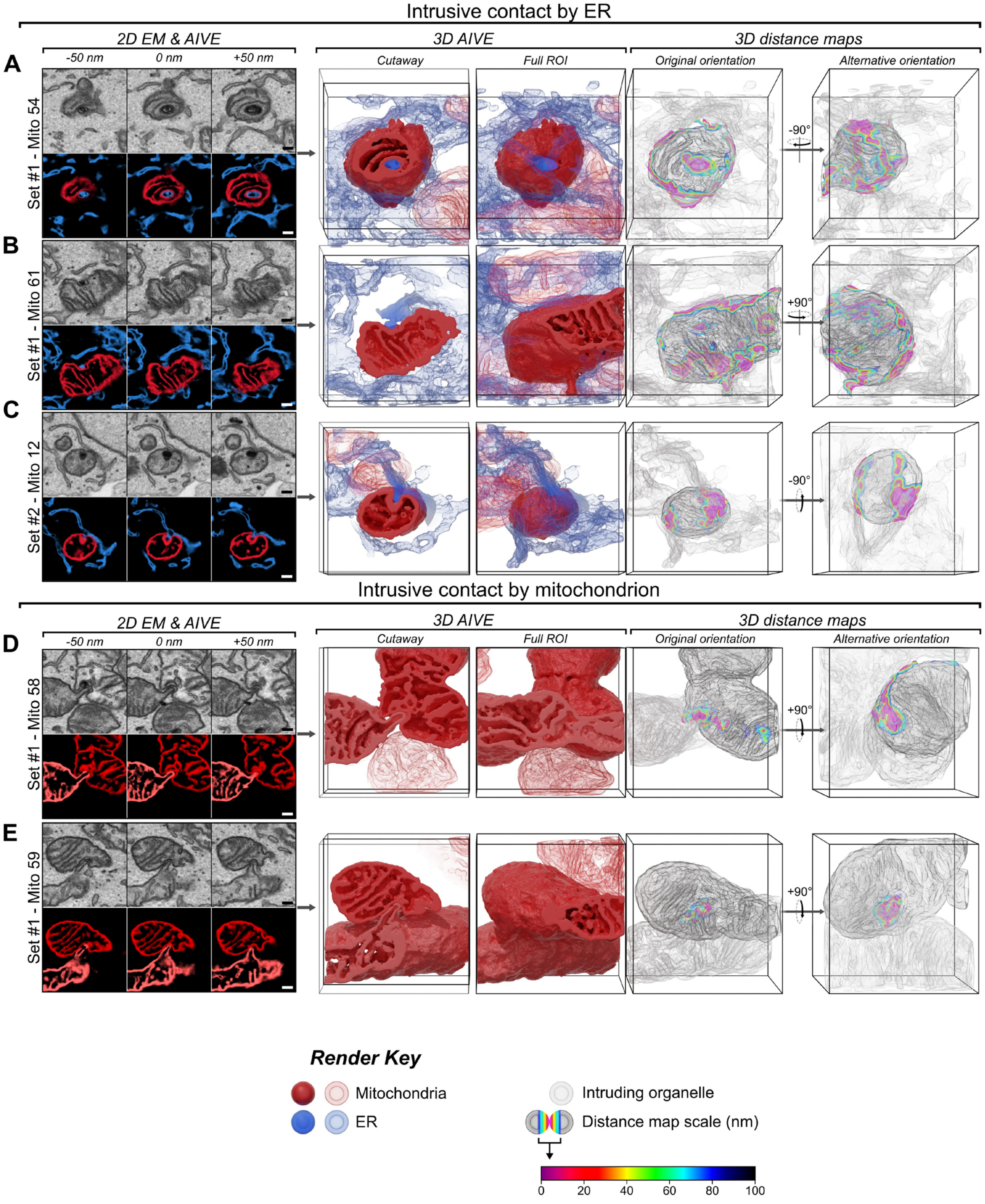
